# Capsaicin activates TRPV1 in human Langerhans cells and inhibits mucosal HIV-1 transmission via secreted CGRP

**DOI:** 10.1101/2021.03.08.434408

**Authors:** Emmanuel Cohen, Aiwei Zhu, Cédric Auffray, Morgane Bomsel, Yonatan Ganor

## Abstract

Upon its mucosal transmission, human immunodeficiency virus type 1 (HIV-1) rapidly targets resident antigen-presenting Langerhans cells (LCs) in genital epithelia, which subsequently trans-infect CD4+ T-cells. We previously described an inhibitory neuro-immune sensory mucosal crosstalk, whereby peripheral pain-sensing nociceptor neurons, innervating all mucosal epithelia and associating with LCs, secret the neuropeptide calcitonin gene-related peptide (CGRP) that strongly inhibits HIV-1 trans-infection. Moreover, we reported that LCs secret low levels of CGRP that are further increased by CGRP itself via an autocrine/paracrine mechanism. As nociceptors secret CGRP following activation of their Ca^2+^ ion channel transient receptor potential vanilloid 1 (TRPV1), we investigated whether LCs also express functional TRPV1. We found that human LCs expressed TRPV1 mRNA and protein. TRPV1 in LCs was functional, as the TRPV1 agonists capsaicin (CP) and resiniferatoxin (RTX) induced Ca^2+^ influx in a dose-dependent manner. Treatment of LCs with CP and the TRPV1 agonist rutaecarpine (Rut) increased CGRP secretion, reaching concentrations close to its IC_50_ for inhibition of HIV-1 trans-infection. Accordingly, CP significantly inhibited HIV-1 trans-infection, which was abrogated by antagonists of both TRPV1 and the CGRP receptor. Finally, pre-treatment of inner foreskin tissue explants with CP markedly increased CGRP secretion, and upon subsequent polarized exposure to HIV-1, inhibited increase in LC-T-cell conjugate formation and T-cell infection. Together, our results reveal that alike nociceptors, LCs express functional TRPV1, whose activation induces CGRP secretion that inhibits mucosal HIV-1 transmission. Our studies could permit re-positioning of formulations containing TRPV1 agonists, already approved for pain relief, as novel topical microbicides against HIV-1.

**Significance Statement:** Upon its sexual transmission, HIV-1 targets different types of mucosal immune cells, such as antigen-presenting Langerhans cells (LCs). In turn, LCs transfer HIV-1 to its principal cellular targets, namely CD4+ T-cells, in a process termed trans-infection. We previously discovered that the mucosal neuropeptide CGRP strongly inhibits trans-infection. CGRP is principally secreted from pain-sensing peripheral neurons termed nociceptors, once activated via their TRPV1 ion channel. Herein, we reveal that LCs also express functional TRPV1, whose activation induces secretion of CGRP that inhibits mucosal HIV-1 transmission. Accordingly, molecules activating TRPV1 and inducing CGRP secretion could be used to prevent mucosal HIV-1 transmission. This approach represents an original neuro-immune strategy to fight HIV-1.

## Introduction

Mucosal tissues evolved complex strategies to detect and protect our body from external threats and invading pathogens via a bi-directional neuro-immune dialogue between sensory peripheral neurons and mucosal immune cells (1). Pain neurons termed nociceptors, innervating all mucosal epithelia, are specialized in sensing noxious stimuli, such as temperatures above 43°C and protons/acidic conditions (2). Their activation is mediated by a large variety of different receptors, of which the non-selective Ca^2+^ ion channel TRPV1 plays a major role (3). A variety of natural compounds also activate TRPV1, for instance capsaicin (CP), the spicy component of chili peppers (4); resiniferatoxin (RTX), a naturally occurring chemical found in the cactus-like plant *Euphorbia resinifera* (5); and rutaecarpine (Rut), an alkaloid isolated from the plant *Evodia rutaecarpa* that is used in traditional Chinese medicine to treat cardiovascular diseases (6). As TRPV1-mediated neuronal excitation evoked by CP, but not other TRPV1 agonists, is followed by a long lasting refractory period with unresponsiveness to additional noxious stimuli (i.e. desensitization), CP topical mucosal formulations and skin patches have been approved clinically as analgesics in painful conditions (7).

Upon TRPV1 activation, nociceptors both conduct pain information into the central nervous system and secret neuropeptides locally at the mucosal level (8). Among these is CGRP (9, 10), a potent vasodilator (11) that contributes to the pathophysiology of several disorders (12). Although nociceptors are the principal source of secreted CGRP upon TRPV1 activation (8), non-neuronal and immune cells also express both molecules (12, 13), e.g. epithelial cells, endothelial cells, fibroblasts, hepatocytes, adipocytes, muscle cells, T-cells and dendritic cells (DCs).

CGRP is a key regulator of inflammatory processes, by mediating neurogenic inflammation (14) and also directly affecting the function of different types of immune cells in a vasodilator-independent manner (15). For instance, nociceptors associate with LCs, and CGRP shifts LCs-mediated antigen presentation and cytokine secretion from Th1 to Th2/Th17 types (15). Such CGRP-mediated bias towards Th2/Th17 immunity is not limited to LCs and occurs also when additional immune cell types are exposed to CGRP, and has both desired and unwanted outcomes during bacterial and fungal infections (16, 17). In contrast, the role of CGRP during viral infections remains largely unexplored.

Mucosal genital epithelia are the principal entry portals of sexually transmitted viruses, including HIV-1 that rapidly targets LCs in both the vagina (18), and inner foreskin as we reported (19–21), followed by transfer of infectious virus from LCs to CD4+ T-cells in a process termed trans-infection (22). Investigating the role of CGRP during mucosal HIV-1 transmission, we discovered a complex neuro-immune interplay whereby CGRP interacts with LCs and modulates a multitude of cellular and molecular processes, resulting in significant inhibition of LCs-mediated HIV-1 trans-infection (23–25). As we reported that LCs secret low basal levels of CGRP that are further increased in an autocrine/paracrine manner by CGRP itself (24), we investigated herein whether LCs express functional TRPV1 whose stimulation could induce CGRP secretion, which in turn would prevent HIV-1 transmission.

## Results

### LCs express functional TRPV1

To test for the presence of TRPV1 in LCs, mRNA was purified from monocyte-derived LCs (MDLCs) and used for qRT-PCR with TRPV1-specific primers. MDLCs prepared from three different healthy individuals all contained TRPV1 mRNA, which was also amplified from total human brain RNA serving as positive control (Fig. 1A).

**Figure 1.**
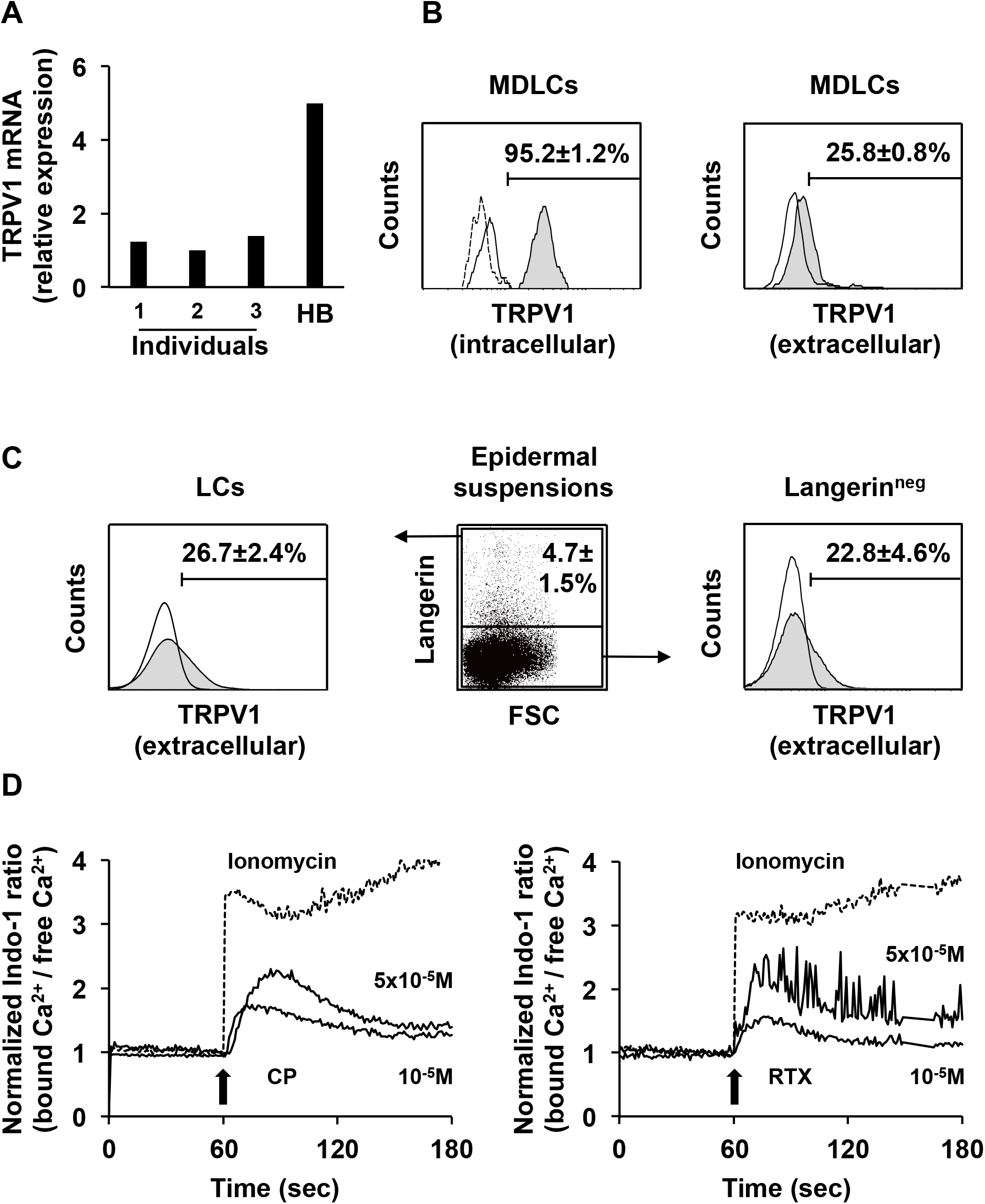
LCs express functional TRPV1. **(A)** Relative TRPV1 mRNA expression in MDLCs from three different human individuals (1–3) normalized to beta actin, with total human brain (HB) mRNA serving as positive control. **(B, C)** MDLCs (B) or epidermal cell suspensions (C) were stained for surface langerin and either intracellular or extracellular TRPV1 expression as indicated, and examined by flow cytometry. Representative overlays show TRPV1 expression (grey histograms) on langerin^+^ gated MDLCs (B) or epidermal cells (C, middle and left), as well as on langerin^neg^ epidermal cells (C, middle and right) vs. matched isotype controls (line histograms). To validate signal specificity, the intracellular TRPV1 Ab was pre-incubated with a blocking peptide before staining (B, left; broken line histogram). Numbers represent mean±SEM percentages of positive cells, derived from n=6 (B, left), n=4 (B, right) and n=3 (C) experiments, using MDLCs or foreskin tissues from different individuals. **(D)** MDLCs were loaded with the Ca^2+^ indicator Indo-1 and examined over time by flow cytometry. Representative graphs (n=3) show the normalized bound/free Ca^2+^ ratio, indicative of Ca^2+^ influx, at baseline (t=0sec, set to 1) and following treatment (t=60sec, arrow) with the indicated concentrations of CP (left) or RTX (right). Ionomycin (1μM; broken lines) served as positive control.

Next, to evaluate TRPV1 expression at the protein level, MDLCs were double stained for surface expression of langerin and either intracellular or extracellular expression of TRPV1, using two different TRPV1 polyclonal antibodies (Abs), and examined by flow cytometry. Langerin surface staining, followed by cell permeabilization and staining with an Ab directed against a cytoplasmic epitope of TRPV1, showed that all langerin^+^ MDLCs expressed TRPV1 intracellularly (Fig. 1B). Signal specificity was validated by pre-incubation of this intracellular TRPV1 Ab with a relevant available blocking peptide, which resulted in loss of staining (Fig. 1B). Staining of MDLCs without permeabilization, using an Ab raised against an epitope including the second extracellular loop of TRPV1, showed that 25% of langerin^+^ MDLCs expressed TRPV1 on their surface (Fig. 1B).

Surface expression of langerin and TRPV1 was also examined in mucosal LCs within epidermal cell suspensions, obtained by enzymatic digestion of inner foreskin tissues using dispase and trypsin (Fig. 1C). Langerin^+^ foreskin LCs accounted for 5% of the cells in such suspensions, and 25% of foreskin LCs expressed TRPV1 on their surface (Fig. 1C). As expected, surface expression of TRPV1 was also detected in 20% of langerin^neg^ foreskin cells (Fig. 1C), comprising primarily of keratinocytes and other immune cells, e.g. T-cells, which express TRPV1 (13). These results further validate the suitability of the extracellular TRPV1 Ab used herein.

To determine whether TRPV1 is functional, MDLCs were loaded with the Ca^2+^ indicator Indo-1, treated with the TRPV1 agonists CP or RTX, and Ca^2+^ influx was measured by flow cytometry. These experiments showed that both TRPV1 agonists induced Ca^2+^ influx in MDLCs in a dose-dependent manner, as did the Ca^2+^ ionophore ionomycin serving as positive control (Fig. 1D). These results show that LCs express surface TRPV1, whose activation by TRPV1 agonists induces Ca^2+^ influx.

### CP inhibits MDLCs-mediated HIV-1 trans-infection via TRPV1 and CGRP

To investigate the potential role of TRPV1 during HIV-1 trans-infection, MDLCs were treated for 24h with either CGRP or the TRPV1 agonist CP. The cells were then pulsed with HIV-1, washed and co-cultured with autologous CD4+ T-cells. A week later, HIV-1 replication was determined by measuring the content of the HIV-1 capsid protein p24 in the co-culture supernatant by ELISA. As we reported (23–25), CGRP strongly inhibited MDLCs-mediated HIV-1 trans-infection in a dose-dependent manner, with maximal inhibition of approximately 90% and IC_50_ at the picomolar range (Fig. 2A). Treatment with CP also inhibited HIV-1 trans-infection in a dose-dependent manner (Fig. 2A). Yet, compared to CGRP, CP had lower efficiency with maximal inhibition of approximately 60%, as well as lower potency with IC_50_ at the micromolar range (Fig. 2A).

**Figure 2.**
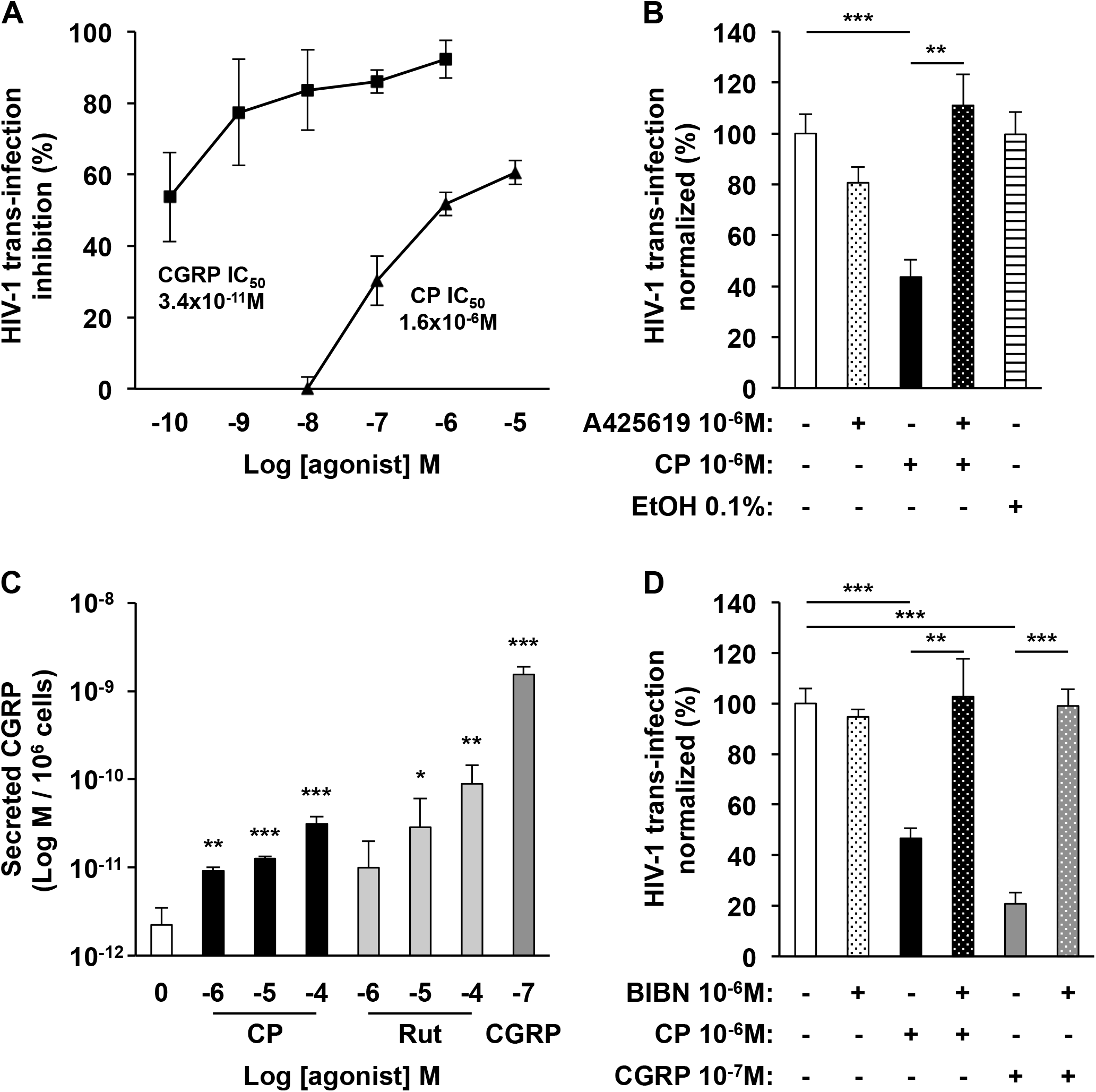
TRPV1 activation in MDLCs inhibits HIV-1 trans-infection via secreted CGRP. **(A, B)** MDLCs were treated for 24h with the indicated molar concentrations of CGRP or CP. The TRPV1 antagonist A425619 was added 15min before addition of CP. The cells were then pulsed with HIV-1 for 4h, washed, and co-cultured with autologous CD4+ T-cells. HIV-1 replication was evaluated a week later by measuring p24 content in the co-culture supernatants using ELISA. In (A), shown are mean±SEM (n=3) percentages of HIV-1 trans-infection inhibition, calculated against untreated cells (i.e. no inhibition). Extrapolated IC_50_ values were 3.4×10^−11^M for CGRP and 1.6×10^−6^M for CP. In (B), shown are mean±SEM (n=4) percentages of HIV-1 trans-infection, normalized against untreated cells serving as the 100% set point. Treatment with 0.1% ethanol (EtOH) served as control. **(C)** MDLCs were treated for 24h with the indicated molar concentrations CGRP, CP or Rut. Culture supernatants were collected immediately after CP and Rut treatment, or following extensive washing and culture in fresh medium for addition 24h after CGRP treatment, and an EIA was used to measure CGRP levels. Shown are mean±SEM (n=3) levels of secreted CGRP per 10^6^ MDLCs. **(D)** MDLCs were treated for 24h with CGRP or CP. The CGRP-R antagonist BIBN4096 (BIBN) was added 15min before addition of agonists. HIV-1 trans-infection was evaluated as above and results (n=4) are shown as in (B). In all graphs, *p<0.0500, **p<0.0050, ***P<0.0005, Student’s t-test.

To confirm that CP-mediated inhibition was mediated by activation of TRPV1, MDLCs were pre-treated with the TRPV1 antagonist A425619 (26) before addition of CP, and HIV-1 trans-infection was determined as above. The TRPV1 antagonist alone had no effect on HIV-1 trans-infection, but completely abrogated CP-mediated inhibition (Fig. 2B). Of note, treatment of MDLCs with 0.1% ethanol, which also activates TRPV1 (27) and was used as diluent for CP, had no effect on HIV-1 trans-infection (Fig. 2B).

To explore the interplay between TRPV1 and CGRP, MDLCs were treated for 24h with the TRPV1 agonists CP and Rut, and the levels of secreted CGRP were next determined in the culture media. Treatment with both TRPV1 agonists induced CGRP secretion from MDLCs in a dose-dependent manner (Fig. 2C), in the range of 0.9-8.9×10^−11^M (Fig. 2C), which spans the IC_50_ of 3.4×10^−11^M for HIV-1 trans-infection inhibition by CGRP (Fig. 2A). For positive control, cells were also treated with exogenous CGRP followed by extensive washing, in order to induce and measure autocrine/paracrine CGRP secretion from MDLCs, as we reported (24). As before, CGRP increased its own secretion from MDLCs, reaching secreted levels at the nanomolar range (Fig. 2C).

Finally, to determine whether CP-induced CGRP secretion is responsible for HIV-1 trans-infection inhibition, MDLCs were pre-treated with the CGRP receptor antagonist BIBN4096 (28) before addition of CGRP or CP. The CGRP receptor antagonist alone had no effect on trans-infection, but completely abrogated both CGRP- and CP-mediated inhibition (Fig. 2D).

These findings show that TRPV1 activation in MDLCs induces CGRP secretion that inhibits HIV-1 trans-infection.

### CP induces CGRP secretion and inhibits HIV-1 transmission in mucosal tissues *ex-vivo*

To extend the *in-vitro* findings above, we tested the activity of CP using mucosal tissues and successive experimental steps (Fig. 3A). Briefly, inner foreskin tissue explants were first submerged in culture media (devoid of CGRP) and left untreated or pre-treated for 24h with CP. The media were then collected and an ELISA was used to measure the levels of secreted CGRP. Next, the explants were transferred to two-chamber transwell inserts and inoculated in a polarized manner for 4h with either non-infected or HIV-1 infected cells, as we previously developed (19–21). The explants were then immediately digested with dispase and trypsin to obtain epidermal cell suspensions. In other experiments, the explants were further incubated for three days submerged in fresh culture medium, and then digested with collagenase and DNase to obtain dermal cell suspensions. The percentages of conjugates between LCs and T-cells, and HIV-1 infection of T-cells, were determined by flow cytometry in either epidermal or dermal suspensions, respectively.

**Figure 3.**
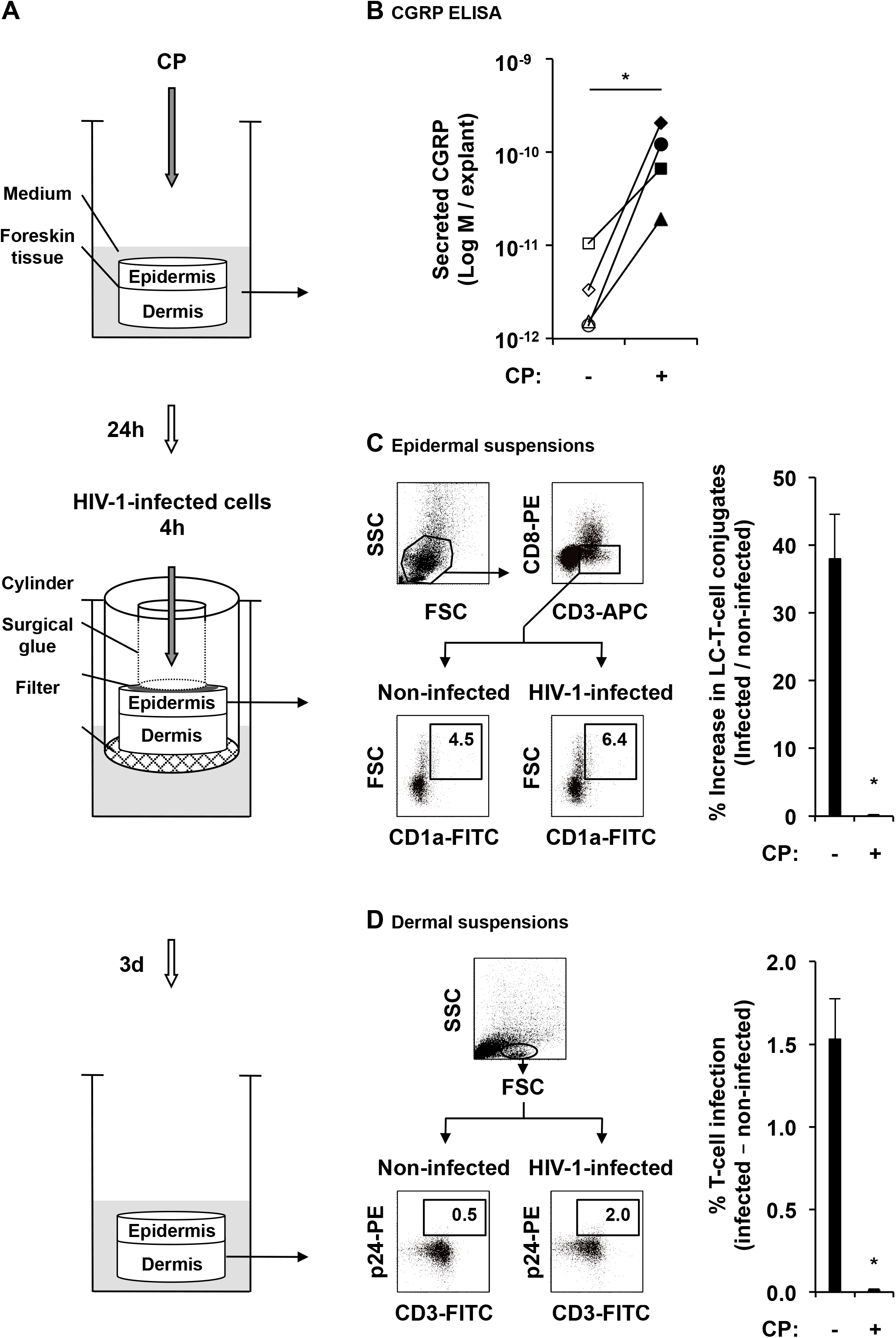
CP inhibits HIV-1 transmission in mucosal inner foreskin tissues *ex-vivo*. **(A)** Schematic representation of the different experimental steps. Inner foreskin tissue explants were submerged in culture media, left untreated or pre-treated with 5×10^−5^M CP, and media were collected 24h later (top). Explants were next washed, transferred to two-chamber transwell inserts and inoculated in a polarized manner with either non-infected or HIV-1-infected cells. Following 4h inoculation, explants were washed and immediately digested with dispase and trypsin to obtain epidermal cell suspensions (middle). Other explants were further incubated for additional three days submerged in fresh culture media, washed and digested with collagenase and DNase to obtain dermal cell suspensions (bottom). **(B)** CGRP levels in 24h explants culture media, measured using CGRP ELISA. Shown are mean values from n=4 explants per condition (for each individual), in matched untreated (open symbols) or CP-treated (closed symbols) explants; *p=0.0286, Mahn-Whitney U-test. **(C)** Representative FACS dot plots of epidermal cell suspensions triple stained for surface expression of CD3, CD8 and CD1a and examined by flow cytometry. After gating out cell debris and FSC^highest^SSC^highest^ keratinocytes (top left), cells were further gated on CD3^+^CD8^−^ T-cells (top right), and the percentages of FSC^high^CD1a^+^ conjugates were determined, following inoculation with either non-infected or HIV-1-infected cells (bottom left and right, respectively). Graph shows mean±SEM (n=3) percentages of increase in conjugate formation, calculated as [(% conjugates following inoculation with HIV-1-infected PBMCs) / (% conjugates following inoculation with non-infected PBMCs) - 1] X 100; *p=0.0380, CP-treated vs. untreated, Student’s t-test. **(D)** Representative FACS dot plots of dermal cell suspensions double stained for surface CD3 and intracellular p24 and examined by flow cytometry. Cells were gated on FSC^low^SSC^low^ lymphocytes (top), and the percentages of CD3+p24^+^ cells were determined, following inoculation with either non-infected or HIV-1-infected PBMCs (bottom left and right, respectively). Graph shows mean±SEM (n=3) percentages of HIV-1-infected T-cells, calculated as [(%CD3+p24+ cells following inoculation with HIV-1-infected PBMCs) - (%CD3+p24+ cells following inoculation with non-infected PBMCs)]; *p=0.0241, CP-treated vs. untreated, Student’s t-test.

In untreated inner foreskin tissue explants, the baseline levels of CGRP secretion were low and ranged from 1.4×10^−12^M to 1.1×10^−11^M, with mean±SEM (n=4) of 4.2±1.9×10^−12^M (Fig. 3B). In all matched explants examined, CP treatment significantly increased CGRP secretion, reaching up to 2.1×10^−10^M, with mean±SEM of 1.2±0.4×10^−10^M, i.e. approximately X30 fold increase (Fig. 3B).

We previously reported that inoculation of inner foreskin tissue explants with HIV-1-infected cells induces secretion of the chemokines CCL5 and thymic stromal lymphopoietin (TSLP) by foreskin keratinocytes, which facilitate increased formation of LC-T-cell conjugates in the epidermis. In turn, LCs locally transfer HIV-1 to T-cells, which migrate to the dermis and become productively infected a few days later (19–21). Pre-treatment with exogenous CGRP inhibits both the increase in LC-T-cell conjugate formation and T-cell infection with HIV-1 (23). Herein, we first confirmed that inoculation with HIV-1-infected cells increased the percentage of high forward scatter (FSC) conjugates between CD1a+ LCs and CD3+CD8- T-cells (Fig. 3C), and resulted in T-cell infection (Fig. 3D). Importantly, CP pre-treatment completely abrogated the increase in conjugate formation (Fig. 3C) and prevented T-cell infection (Fig. 3D), induced by inoculation of HIV-1-infected cells. These findings show that CP induces CGRP secretion and inhibits HIV-1 transmission in mucosal tissues *ex-vivo*.

## Discussion

We report herein on a novel neuro-immune mechanism that limits mucosal HIV-1 transmission. Accordingly, CP treatment of isolated LCs or inner foreskin tissues induces activation of TRPV1 and subsequent secretion of CGRP, which in turn inhibits LCs-mediated HIV-1 trans-infection *in-vitro* and HIV-1 transmission *ex-vivo*.

TRPV1 immunoreactivity in human skin LCs was previously reported by one study (using a polyclonal goat TRPV1 Ab that is no longer available commercially) (29), but was not confirmed by another study (using a custom-produced polyclonal rabbit TRPV1 Ab) (30). We used both qRT-PCR and immunolabeling followed by flow cytometry, to show that MDLCs and foreskin LCs express TRPV1 mRNA and protein. Interestingly, while all langerin^+^ MDLCs express TRPV1 intracellularly, only a subset of MDLCs and foreskin LCs cells showed surface expression of TRPV1. Such discrepancy might be related to differential epitope accessibility of the two TRPV1 Abs we used. The intracellular TRPV1 Ab is directed against an epitope located within the cytoplasmic C-terminus region of TRPV1, which would be readily accessible to Ab binding upon permeabilization. In contrast, the extracellular TRPV1 epitope is directed against an epitope spanning the second extracellular loop and part of the fourth transmembrane domain of TRPV1, and might be less accessible. Alternatively, these results might suggest the LCs contain an intracellular pool of TRPV1, which is only partially transported to the plasma membrane. Indeed, TRPV1 activity is positively regulated by different factors that promote TRPV1 surface localization (3). Moreover, TRPV1 activation leads to its desensitization that is mediated by both agonist-induced conformational changes at the channel level (31), as well as by modification in TRPV1 recycling (32). While our results show that LCs express functional surface TRPV1 that facilitates Ca^2+^ influx upon activation by both CP and RTX, the exact mechanisms mediating TRPV1 transport, recycling, and potentially desensitization, in LCs remain to be determined.

CGRP is expressed by approximately 45% of dorsal root ganglia (DRG) neurons, predominantly in small diameter unmyelinated C-fibers, as well as small-to-medium diameter Aδ fibers (33). About half of these CGRP-expressing nociceptors, which innervate all mucosal epithelia including the foreskin, also express TRPV1 and respond to noxious stimuli and natural TRPV1 agonists by secreting CGRP (34). We show that a similar sensory mechanism operates in LCs, which express TRPV1 and secret CGRP following TRPV1 activation. These results are the first to demonstrate a functional role for TRPV1 in LCs, and extend previous observations of a similar interplay between TRPV1 and CGRP in other non-neuronal cells, e.g. CGRP release has been linked to TRPV1 activation in keratinocytes (35) and DCs (36).

Our measurements of CGRP secretion following treatment with 10^−5^M CP indicate that 1×10^6^ isolated MDLCs secrete 1.3×10^−11^M CGRP (Fig. 2C), while treatment with 5×10^−5^M CP of inner foreskin tissue explants induces secretion of 1.2×10^−10^M CGRP (Fig. 3B). Based on our previous quantitative analysis of LCs density in 4μm tissue sections of the inner foreskin (≈600 LCs / mm^2^ (19, 20)) and an average thickness of 0.1mm of human skin epidermis, we estimate that ≈0.8×10^6^ LCs will be present in each of the inner foreskin tissue explants (8mm in diameter). Hence, resident LCs within the tissue explants could contribute approximately 1/10 of total secreted CGRP upon CP activation. These results suggest that LCs are an important, yet not the principal source of mucosal CGRP. We speculate that although nociceptors are sectioned during tissue sampling, they might still maintain the capacity to respond to TRPV1 agonists by secreting CGRP, which is pre-stored in their mucosal nerve terminals. In addition, foreskin keratinocytes could represent another source of CGRP, which can be secreted from these cells upon TRPV1 activation by CP (35). Accordingly, keratinocytes were suggested to function as unconventional (non-neuronal) sensors of a variety of environmental factors that communicate with nociceptors, potentially representing a key part of the sensory system of the skin (37). In accordance, a recent study reported that TRPV1 activation in keratinocytes is sufficient to evoke acute nociception-related responses (38). It would be instrumental now to determine the exact contribution of nociceptors and keratinocytes to CP-induced mucosal CGRP secretion, test the expression of other TRP channels in human LCs, and determine whether and how LCs participate in the sensory network in the skin and other mucosal epithelia.

Although T-cells are CGRP-responsive (39), our previous studies showed that pre-treatment of CD4+ T-cells with CGRP, rather than LCs, has no effect on HIV-1 trans-infection (23). Similarly, our recent *in-vivo* study suggests that CGRP acts by targeting LCs and delaying HIV-1 dissemination to CD4+ T-cells in humanized BLT mice (40). These findings indicate that exogenous CGRP and/or endogenous CGRP (secreted from LCs upon TRPV1 activation, as reported herein) probably act on LCs and not T-cells, to inhibit HIV-1 trans-infection. Yet, CD4+ T-cells express TRPV1 whose levels increase upon T-cell activation (41) and are CP-responsive (41, 42), suggesting that CP could interfere with their HIV-1 infection. Our on-going studies are now aimed at defining CGRP-dependent/independent mechanisms, via which CP could inhibit directly HIV-1 infection of CD4+ T-cells.

Formulations containing CP are clinically approved for local pain control. For instance, topical CP is used to treat HIV-1-associated distal sensory polyneuropathy and postherpetic neuralgia (7), and CP creams are applied vaginally to treat vulvar vestibulitis syndrome (43). As CP is a known irritant, its topical application induces an initial burning ‘flare’ sensation associated with nociceptor activation. Such discomfort is limited to the site of application and resolves after the first few days (7). On-going studies are exploring strategies to improve CP bioavailability (e.g. CP nanoparticles), which may decrease CP dosage, leading to minimal side effects (44).

Together, our study provides ‘proof-of-concept’ and support for the clinical use of formulations containing CGRP and/or TRPV1 agonists as novel topical microbicides active against HIV-1. This neuro-immune based approach offers an original and potentially advantageous alternative for prevention of HIV-1 transmission.

## Materials and Methods

### Cells and tissues

Peripheral blood mononuclear cells (PBMCs) from healthy HIV-1 seronegative individuals were separated from whole blood by standard Ficoll gradient. CD14+ monocytes and CD4+ T-cells were purified from PBMCs by negative magnetic selection (Stemcell Technologies, Grenoble, France). Monocytes (1×10^6^/well, in 12-wells plate) were differentiated into MDLCs in complete medium (RPMI 1640 medium, 10% fetal bovine serum (FBS), 2mM glutamine, 100U/ml penicillin and 100mg/ml streptomycin (Gibco Invitrogen, Carlsbad, CA)) supplemented with 100ng/ml granulocyte-macrophage colony-stimulating factor (GM-CSF), 10ng/ml interleukin 4 (IL-4) and 10ng/ml transforming growth factor beta 1 (TGFβ1) (R&D systems, Minneapolis, MN), and used between days 7-9 of differentiation.

Normal foreskin tissues were obtained from healthy adults undergoing circumcision (Urology Service, Cochin Hospital, Paris), under informed consent and ethical approval (Comités de Protection des Personnes CPP Paris-IdF XI, N.11016). Inner foreskins (distinguished by their lighter color and morphology) were separated mechanically, and remaining fat and muscle tissue was removed from the dermal side. Epidermal cell suspension were prepared as we described (19, 20). Briefly, tissue pieces were incubated with their epidermal side facing up in RPMI 1640 medium supplemented with 2.4 U/ml Dispase II (Roche Diagnostics GmbH, Mannheim, Germany) overnight at 4°C. The epidermis and dermis were then mechanically separated using forceps and epidermal cell suspensions were prepared by incubating the epidermal sheets in 0.05% Trypsin/EDTA (Gibco) for 10 min at 37°C, followed by inactivation of trypsin with FBS, mechanical disruption using a 10ml pipette, filtration of released cells through 100μm nylon cell strainers and centrifugation.

### Virus and infected cells

Viral stocks of the HIV-1 molecular clone ADA and the primary isolate 93BR029 (both clade B, R5 tropic; NIH AIDS reagent program) were prepared by transfection of 293T cells or by amplification on phytohaemagglutinin (PHA)/IL-2-stimulated PBMCs, respectively, and quantified using the p24 Innotest HIV-1 ELISA (Fujirebio, Courtaboeuf, France). HIV-1 V29-infected PBMCs were prepared as we reported (19).

### TRPV1 qRT-PCR

RNA purification from MDLCs and cDNA reverse transcription were performed as we previously described (24). Total human brain (HB) RNA (Clontec, Mountain View, CA, USA) served as positive control. qRT-PCR was performed with the TRPV1 TaqMan Gene Expression Assay (TheroFisher Scientific, Waltham, MA USA). The 20μl PCR reaction mixture contained 2μl of cDNA prepared from 1μg total RNA, 10μl of Master Mix 2X from TaqMan RNA-to-Ct *1-step* Kit (TheroFisher Scientific), and 1μl of each primer. Reactions were performed in triplicates, with beta actin as the internal control. Thermocycler conditions consisted of pre-incubation at 95°C for 10min for 1 cycle, followed by 45 cycles of denaturation at 95°C for 15sec and annealing/extension at 60°C for 60sec. Amplification, data acquisition and analysis were carried out using the LightCycler 480 Software (Roche, Mannheim, Germany) and relative expression levels were quantified using the 2^−ΔCt^ method.

### TRPV1 flow cytometry

MDLCs (1×10^5^/well) or epidermal cells (5×10^5^/well) were stained on ice for 30min with 1:20 dilution of TRPV1 rabbit polyclonal Ab raised against an extracellular epitope of human TRPV1 (amino acids 531-541; TheroFisher Scientific, Waltham, MA USA), followed by 1:200 dilution of secondary Cy5-conjugated donkey-anti-rabbit-IgG (Jackson Immunoresearch, West Grove, PA, USA) and 10μl of PE-conjugated mouse-anti-human langerin Ab (Clone DCGM4; Beckman Coulter, Marseille, France). In other experiments, MDLCs were first stained on ice for 30min with the langerin Ab, fixed, permeabilized, and stained at room temperature for 30min with 8.5μl/ml of TRPV1 rabbit polyclonal Ab directed against an intracellular epitope of rat TRPV1 (amino acids 824-838; Alomone, Jerusalem, Israel), followed by 1:100 dilution of secondary FITC-conjugated goat-anti-rabbit-IgG (Jackson). For blocking experiments, the intracellular TRPV1 Ab was pre-incubated at room temperature with 1:1 ratio of its associated blocking peptide before staining. Cells stained with matched isotype Abs served as negative control. Fluorescent profiles were recorded using a Guava easyCyte flow cytometer and analyzed with InCyte software (Merck Millipore, Guyancourt, France).

### Intracellular Ca^2+^

MDLCs (4×10^5^/tube) were loaded for 30min at 37°C with 1μM of the membrane-permeable fluorescent Ca^2+^ indicator dye Indo-1 AM, washed, equilibrated for 5min at 37°C before acquisition, and Ca^2+^ traces were recorded by flow cytometry using an LSR-II (BD Biosciences), as we reported (45). After the establishment of a stable baseline for 60sec, cells were stimulated with the indicated concentrations of CP, RTX or ionomycin (Sigma, St. Louis, MO, USA), and traces were recorded for additional 120sec. Data was analyzed with Diva software (BD Biosciences).

### HIV-1 trans-infection

MDLCs (1×10^5^/well) were treated for 24h at 37°C with the indicated concentrations of CGRP, CP or absolute ethanol (Sigma) in a 96 round-bottom wells plate (200μl/well final). The TRPV1 antagonist A425619 (Tocris / Biotechne, Minneapolis, MN, USA) and the CGRP receptor anatagonist BIBN4096 (Sigma) were added 15min before agonists. The cells were then washed, pulsed with HIV-1 ADA (1ng p24 corresponding to multiplicity of infection (MOI) of 0.2) for 4h, whashed again, and incubated with autologous CD4+ T-cells (3×10^5^/well). HIV-1 p24 content was measured a week later in the co-culture supernatants using p24 ELISA (Fujirebio, Courtaboeuf, France). IC_50_ values were calculated with Prism software (GraphPad) using the log(inhibitor) vs. normalized response - variable slope model.

### Inner foreskin tissue explants

Round inner foreskin tissue pieces were cut using a 8mm diameter Harris Uni-Core, transferred to 24-wells plate and incubated submerged for 24h at 37°C in 1ml coplete RPMI medium, alone or supplemented with 5×10^−5^M CP (four explants per condition). The media were next collected and tested for CGRP content (see below). Next, the tissues were washed, transferred to two-chamber transwell inserts (Sigma), and inoculated in a polarized manner for 4h at 37°c with either non-infected or HIV-1 infected PBMCs (in duplicates), as we described (19, 20). Epidermal cell suspensions were prepared immediately after inoculation, as described above. Pooled cells of each duplicate were resuspended in phospahte-buffered saline (PBS), transferred to a 96 round-bottom wells plate and stained for 30min on ice with 10μl of FITC-conjugated mouse-anti-human CD1a, PE-conjugated mouse-anti-human CD8 and APC-conjugated mouse-anti-human CD3 Abs (BD Pharmingen, San Jose, CA), diluted in PBS to a final volume of 50μl/well. Dermal cell suspensions were prepared following washing of the explants, additional incubation for three days at 37°C submerged in 1ml fresh medium, and subsequent enzymatic digestion with collagenase and DNase, as we described (19, 20). Cells were surface stained as above using FITC-conjugated mouse-anti-human CD3 Ab (Pharmingen), fixed, permeabilized, and stained for 30min at room temperature with 1:160 dilution of PE-conjugated mouse-anti-human Ab to HIV-1 p24 and core antigens (Beckman Coulter, Fullerton, CA). Fluorescent profiles were recorded using a Guava easyCyte and InCyte software.

### CGRP secretion

MDLCs (5×10^5^/well) were treated for 24h at 37°c with the indicated concentrations of CGRP, CP or Rut in a 96 round-bottom wells plate (200μl/well final). The culture supernatants were collected immediately following CP and Rut treatment or following extensive washing and incubation for additional 24h at 37°C in fresh medium following CGRP treatment. CGRP levels were determined using a competitive CGRP enzyme immunoassay (EIA; Peninsula, San Carlos, CA). CGRP secretion from inner foreskin tissue explants into the culture media was determined using a direct CGRP ELISA (Bertin, Rockville, MD).

### Statistical analysis

Statistical significance was analyzed with the two-tailed Student’s t-test. For comparing secreted CGRP levels from matched tissue explants, pair-wise comparisons were performed with the non-parametric Mann-Whitney U-test.

## Acknowledgments

E.C. was supported by a PhD fellowship from the French national agency for HIV-1 and hepatitis research (ANRS), A.Z. was supported by a PhD fellowship from the Chinese scientific council, and the study was funded by a grant to Y.G. from SIDACTION (Aide aux Equipes, Ref 17-2-AEQ-11613).

## References

1 Talbot S, Foster SL, & Woolf CJ (2016) Neuroimmunity: Physiology and Pathology. Annu Rev Immunol 34:421–447.

2 Dubin AE & Patapoutian A (2010) Nociceptors: the sensors of the pain pathway. J Clin Invest 120(11):3760–3772.

3 Nilius B & Szallasi A (2014) Transient receptor potential channels as drug targets: from the science of basic research to the art of medicine. Pharmacol Rev 66(3):676–814.

4 Caterina MJ, et al. (1997) The capsaicin receptor: a heat-activated ion channel in the pain pathway. Nature 389(6653):816–824.

5 Szallasi A & Blumberg PM (1989) Resiniferatoxin, a phorbol-related diterpene, acts as an ultrapotent analog of capsaicin, the irritant constituent in red pepper. Neuroscience 30(2):515–520.

6 Jia S & Hu C (2010) Pharmacological effects of rutaecarpine as a cardiovascular protective agent. Molecules 15(3):1873–1881.

7 Hall OM, et al. (2020) Novel Agents in Neuropathic Pain, the Role of Capsaicin: Pharmacology, Efficacy, Side Effects, Different Preparations. Curr Pain Headache Rep 24(9):53.

8 De Logu F, Nassini R, Landini L, & Geppetti P (2019) Pathways of CGRP Release from Primary Sensory Neurons. Handb Exp Pharmacol 255:65–84.

9 Amara SG, Jonas V, Rosenfeld MG, Ong ES, & Evans RM (1982) Alternative RNA processing in calcitonin gene expression generates mRNAs encoding different polypeptide products. Nature 298(5871):240–244.

10 Rosenfeld MG, et al. (1983) Production of a novel neuropeptide encoded by the calcitonin gene via tissue-specific RNA processing. Nature 304(5922):129–135.

11 Brain SD, Williams TJ, Tippins JR, Morris HR, & MacIntyre I (1985) Calcitonin gene-related peptide is a potent vasodilator. Nature 313(5997):54–56.

12 Russell FA, King R, Smillie SJ, Kodji X, & Brain SD (2014) Calcitonin gene-related peptide: physiology and pathophysiology. Physiol Rev 94(4):1099–1142.

13 Gunthorpe MJ & Szallasi A (2008) Peripheral TRPV1 receptors as targets for drug development: new molecules and mechanisms. Curr Pharm Des 14(1):32–41.

14 Sousa-Valente J & Brain SD (2018) A historical perspective on the role of sensory nerves in neurogenic inflammation. Semin Immunopathol 40(3):229–236.

15 Granstein RD, Wagner JA, Stohl LL, & Ding W (2015) Calcitonin gene-related peptide: key regulator of cutaneous immunity. Acta physiologica 213(3):586–594.

16 Baral P, Udit S, & Chiu IM (2019) Pain and immunity: implications for host defence. Nat Rev Immunol 19(7):433–447.

17 Pinho-Ribeiro FA, Verri WA, Jr., & Chiu IM (2017) Nociceptor Sensory Neuron-Immune Interactions in Pain and Inflammation. Trends Immunol 38(1):5–19.

18 Hladik F, et al. (2007) Initial events in establishing vaginal entry and infection by human immunodeficiency virus type-1. Immunity 26(2):257–270.

19 Ganor Y, et al. (2010) Within 1 h, HIV-1 uses viral synapses to enter efficiently the inner, but not outer, foreskin mucosa and engages Langerhans-T cell conjugates. Mucosal Immunol 3(5):506–522.

20 Zhou Z, et al. (2011) HIV-1 Efficient Entry in Inner Foreskin Is Mediated by Elevated CCL5/RANTES that Recruits T Cells and Fuels Conjugate Formation with Langerhans Cells. PLoS Pathog 7(6):e1002100.

21 Zhou Z, et al. (2018) The HIV-1 viral synapse signals human foreskin keratinocytes to secrete thymic stromal lymphopoietin facilitating HIV-1 foreskin entry. Mucosal Immunol 11(1):158–171.

22 Nasr N, et al. (2014) Inhibition of two temporal phases of HIV-1 transfer from primary Langerhans cells to T cells: the role of langerin. J Immunol 193(5):2554–2564.

23 Ganor Y, et al. (2013) Calcitonin gene-related peptide inhibits Langerhans cell-mediated HIV-1 transmission. J Exp Med 210(11):2161–2170.

24 Ganor Y, Drillet-Dangeard AS, & Bomsel M (2015) Calcitonin gene-related peptide inhibits human immunodeficiency type 1 transmission by Langerhans cells via an autocrine/paracrine feedback mechanism. Acta physiologica 213(2):432–441.

25 Bomsel M & Ganor Y (2017) Calcitonin Gene-Related Peptide Induces HIV-1 Proteasomal Degradation in Mucosal Langerhans Cells. J Virol 91(23): e01205–17.

26 McGaraughty S, Chu KL, Faltynek CR, & Jarvis MF (2006) Systemic and site-specific effects of A-425619, a selective TRPV1 receptor antagonist, on wide dynamic range neurons in CFA-treated and uninjured rats. J Neurophysiol 95(1):18–25.

27 Trevisani M, et al. (2002) Ethanol elicits and potentiates nociceptor responses via the vanilloid receptor-1. Nat Neurosci 5(6):546–551.

28 Doods H, et al. (2000) Pharmacological profile of BIBN4096BS, the first selective small molecule CGRP antagonist. Br J Pharmacol 129(3):420–423.

29 Bodo E, et al. (2004) Vanilloid receptor-1 (VR1) is widely expressed on various epithelial and mesenchymal cell types of human skin. J Invest Dermatol 123(2):410–413.

30 Stander S, et al. (2004) Expression of vanilloid receptor subtype 1 in cutaneous sensory nerve fibers, mast cells, and epithelial cells of appendage structures. Exp Dermatol 13(3):129–139.

31 Touska F, Marsakova L, Teisinger J, & Vlachova V (2011) A “cute” desensitization of TRPV1. Curr Pharm Biotechnol 12(1):122–129.

32 Tian Q, et al. (2019) Recovery from tachyphylaxis of TRPV1 coincides with recycling to the surface membrane. Proc Natl Acad Sci U S A 116(11):5170–5175.

33 Iyengar S, Ossipov MH, & Johnson KW (2017) The role of calcitonin gene-related peptide in peripheral and central pain mechanisms including migraine. Pain 158(4):543–559.

34 McCoy ES, Taylor-Blake B, & Zylka MJ (2012) CGRPalpha-expressing sensory neurons respond to stimuli that evoke sensations of pain and itch. PLoS One 7(5):e36355.

35 Zhang R, et al. (2018) Sirtuin6 inhibits c-triggered inflammation through TLR4 abrogation regulated by ROS and TRPV1/CGRP. J Cell Biochem 119(11):9141–9153.

36 Assas BM, Wakid MH, Zakai HA, Miyan JA, & Pennock JL (2016) Transient receptor potential vanilloid 1 expression and function in splenic dendritic cells: a potential role in immune homeostasis. Immunology 147(3):292–304.

37 Moran MM, McAlexander MA, Biro T, & Szallasi A (2011) Transient receptor potential channels as therapeutic targets. Nat Rev Drug Discov 10(8):601–620.

38 Pang Z, et al. (2015) Selective keratinocyte stimulation is sufficient to evoke nociception in mice. Pain 156(4):656–665.

39 Assas BM, Pennock JI, & Miyan JA (2014) Calcitonin gene-related peptide is a key neurotransmitter in the neuro-immune axis. Front Neurosci 8:23.

40 Bomsel M, et al. (2020) Full-length native CGRP neuropeptide and its stable analogue SAX, but not CGRP peptide fragments, inhibit mucosal HIV-1 transmission. Under Review.

41 Majhi RK, et al. (2015) Functional expression of TRPV channels in T cells and their implications in immune regulation. FEBS J 282(14):2661–2681.

42 Bertin S, et al. (2014) The ion channel TRPV1 regulates the activation and proinflammatory properties of CD4(+) T cells. Nat Immunol 15(11):1055–1063.

43 Steinberg AC, Oyama IA, Rejba AE, Kellogg-Spadt S, & Whitmore KE (2005) Capsaicin for the treatment of vulvar vestibulitis. Am J Obstet Gynecol 192(5):1549–1553.

44 Rollyson WD, et al. (2014) Bioavailability of capsaicin and its implications for drug delivery. J Control Release 196:96–105.

45 Guichard V, et al. (2017) Calcium-mediated shaping of naive CD4 T-cell phenotype and function. Elife 6.

